# Sequence variation of rare outer membrane protein β-barrel domains in clinical strains provides insights into the evolution of *Treponema pallidum* subsp. *pallidum*, the syphilis spirochete

**DOI:** 10.1101/318097

**Authors:** Sanjiv Kumar, Melissa J. Caimano, Arvind Anand, Abhishek Dey, Kelly L. Hawley, Morgan E. LeDoyt, Carson J. La Vake, Adriana R. Cruz, Lady G. Ramirez, Lenka Paštěková, Irina Bezsonova, David Šmajs, Juan C. Salazar, Justin D. Radolf

## Abstract

In recent years, considerable progress has been made in topologically and functionally characterizing integral outer membrane proteins (OMPs) of *Treponema pallidum* subspecies *pallidum* (*TPA*), the syphilis spirochete, and identifying its surface-exposed β-barrel domains. Extracellular loops in OMPs of Gram-negative bacteria are known to be highly variable. We examined the sequence diversity of β-barrel-encoding regions of *tprC*, *tprD*, and *bamA*, in 31 specimens from Cali, Colombia; San Francisco, California; and the Czech Republic and compared them to allelic variants in the 41 reference genomes in the NCBI database. To establish a phylogenetic framework, we used *tp0548* genotyping and *tp0558* sequences to assign strains to the Nichols or SS14 clades. We found that (i) β-barrels in clinical strains could be grouped according to allelic variants in *TPA* reference genomes; (ii) for all three OMP loci, clinical strains within the Nichols or SS14 clades often harbored β-barrel variants that differed from the Nichols and SS14 reference strains; and (iii) OMP variable regions often reside in predicted extracellular loops containing B-cell epitopes. Based upon structural models, non-conservative amino acid substitutions in predicted transmembrane β-strands of TprC and TprD2 could give rise to functional differences in their porin channels. OMP profiles of some clinical strains were mosaics of different reference strains and did not correlate with results from enhanced molecular typing. Our observations suggest that human host selection pressures drive *TPA* OMP diversity and that genetic exchange contributes to the evolutionary biology of *TPA*. They also set the stage for topology-based analysis of antibody responses against OMPs and help frame strategies for syphilis vaccine development.

**IMPORTANCE:** Despite recent progress characterizing outer membrane proteins (OMPs) of *Treponema pallidum* (*TPA*), little is known about how their surface-exposed, β-barrel-forming domains vary among strains circulating within high-risk populations. In this study, sequences for the β-barrel-encoding regions of three OMP loci, *tprC*, *tprD*, and *bamA,* in *TPA* from a large number of patient specimens from geographically disparate sites were examined. Structural models predict that sequence variation within β-barrel domains occurred predominantly within predicted extracellular loops. Amino acid substitutions in predicted transmembrane strands that could potentially affect porin channel function also were noted. Our findings suggest that selection pressures exerted by human populations drive *TPA* OMP diversity and that recombination at OMP loci contributes to the evolutionary biology of syphilis spirochetes. These results also set the stage for topology-based analysis of antibody responses that promote clearance of *TPA* and frame strategies for vaccine development based upon conserved OMP extracellular loops.

## INTRODUCTION

After years of steady decline during the 1990s, syphilis, a sexually transmitted infection caused by the uncultivatable spirochete *Treponema pallidum* subsp. *pallidum* (*TPA*), has undergone a dramatic resurgence in the United States, particularly among men who have sex with men (1). Syphilis also poses a major threat globally with an estimated 5.6 million new cases annually and 350,000 adverse pregnancy outcomes due to mother-to-child transmission (2). The failure of epidemiological approaches to curtail the spread of syphilis underscores the need for a vaccine capable of inducing protective antibody responses against geographically-widespread and genetically-diverse *TPA* strains (3, 4). *TPA* has been designated “the stealth pathogen” based on its ability to evade innate and adaptive immune responses for protracted periods, permitting repeated bouts of hematogenous dissemination and invasion of numerous organs, including the central nervous system and the fetal-placental barrier during pregnancy (5-7). While the appearance of opsonic antibodies is widely regarded as a turning point in the battle between host and pathogen (8, 9), the targets of antibodies that promote bacterial clearance during syphilitic infection are largely unidentified. How immune pressures in high-risk populations influence the epidemiology of syphilis and the evolutionary biology of *TPA* also is poorly understood.

Dual membrane bacteria have evolved a unique class of integral outer membrane protein in which anti-parallel, amphipathic β-strands circularize to form a closed barrel structure, often creating a central aqueous channel that permits uptake of nutrients and efflux of waste products (10-12). Extracellular loops bridge adjacent transmembrane strands, extending from the OM into the external milieu (13). Protective B-cell determinants reside in the extracellular loops and undergo sequence/antigenic variation to circumvent herd immunity against previously circulating strains (14-18). Extracellular loops also play critical roles in disease pathogenesis by promoting interactions with host cells and tissue components and protecting the bacterium against innate clearance mechanisms, such as complement-mediated lysis and neutrophil engulfment (19-25).

Multiple factors have impeded efforts to identify the syphilis spirochete’s integral OMPs. These include the recalcitrance of *TPA* to *in vitro* cultivation (26, 27), the fragility of its outer membrane (28, 29), its relatively low abundance of outer membrane-spanning proteins (30, 31), and the lack of strong sequence relatedness between *TPA* OMPs and well-characterized proteins in Gram-negative OMs (32, 33). To circumvent these obstacles, computational methods were employed to mine the *TPA* Nichols strain genome for proteins predicted to form OM-associated β-barrels. This bioinformatics approach, combined with a battery of biophysical and cellular localization techniques, including opsonophagocytosis assays, yielded a panel of candidate OMPs (33-37). One of these, TP0326/BamA, is the central component of the molecular machine that chaperones newly exported precursor OMPs from the periplasm into the OM (37, 38). A homology model based on the solved structure of the *Neisseria gonorrhoeae* ortholog (38) predicts that the β-barrel of *TPA* BamA contains 16 transmembrane β-strands and eight extracellular loops (37). One extracellular loop, L4, was previously shown using human syphilitic sera to contain an immunodominant epitope; antibodies against this loop also promoted opsonization of *TPA* by rabbit peritoneal macrophages (37). A second group of candidate OMPs, TprC/D (TP0117/TP0131) and TprI (TP0620), are members of the paralogous *T. pallidum* repeat (Tpr) family (39, 40). Analysis of recombinant TprC/D and TprI suggests that their β-barrel domains form aqueous channels in liposomes (35, 36), which is consistent with their potential functions as porins. As with porins from Gram-negative bacteria (11), TprC/D and TprI form trimers, with the β-barrel domains being essential for trimerization (35, 36). A similar bipartite topology with a periplasmic N-terminal (MOSP^N^) and OM-embedded C-terminal (MOSP^C^) trimeric β-barrel also has been demonstrated for the major outer sheath protein (MOSP) of the oral commensal *T. denticola,* the parental ortholog for the Tpr family (41, 42). Surface epitope mapping of subfamily I members based on accurate structural models has yet to be performed.

Previously, Centurion-Lara and co-workers (40) performed a detailed analysis of *tpr* genes for a small number of isolates from four subspecies of pathogenic treponemes. Their predictions regarding sequence variability and immune pressure, however, were based on structural models for TprC/D and TprI that used the full-length polypeptides rather than just the OM-embedded β-barrel-forming MOSP^C^ domains. Thus, no study to date has looked at sequence variation within the regions of *TPA* OMPs known to reside at the host-pathogen-interface (*i.e*., surface-exposed) in multiple *TPA* strains circulating within at-risk populations. Herein, we examined the β-barrel-encoding (MOSP^C^) domains of the *tprC*, *tprD*, and *bamA* in DNA extracted from *TPA* within 31 clinical specimens obtained from early syphilis patients in Cali, Colombia (43, 44), San Francisco, CA (45), and the Czech Republic (46, 47). The resulting sequences were compared to the corresponding loci within the 41 *TPA* reference genomes available from NCBI databases. Based on *s*tructural models for TprC/D and BamA, much of the sequence variability within all three OMPs is predicted to lie within extracellular loops containing B-cell epitopes. The models also identified amino acid substitutions in predicted transmembrane β-strands for TprC/D and TprD2, which could affect the selectivity of their porin channels. Lastly, OMP profiles of clinical strains at all three loci appear to be mosaics of alleles represented in the *TPA* reference genomes and, importantly, did not correlate with results from enhanced molecular typing. The findings presented herein are consistent with the notion that selection pressures within human populations drive *TPA* OMP diversity and that genetic exchange within and between the Nichols and SS14 clades contributes to the evolutionary biology of syphilis spirochetes. These results also set the stage for topology-based analyses of antibody responses that promote clearance of individual *TPA* strains and, importantly, could be used to develop a broadly protective vaccine based on conserved extracellular loops.

## RESULTS

### Patients and clinical samples

Skin biopsies from secondary syphilis rashes were obtained from patients seen in Cali, Colombia; swabs from exudative lesions were obtained from early syphilis patients in San Francisco, CA (SF) and in Brno and Prague, Czech Republic (CZ). Specimens from Cali and SF were chosen for amplification and sequencing of β-barrel regions based on treponemal burdens determined by *polA* qPCR. CZ specimens selected were PCR-positive for all typing loci tested (47), reflecting high *TPA* burdens. Table S1 contains a summary of available demographic information for the samples used in this study.

### *TPA* in clinical samples belong to the Nichols and SS14 clades

Prior multilocus sequence analyses demonstrated that syphilis spirochetes cluster into two taxonomic groups, or clades, arbitrarily named after the Nichols and SS14 reference strains (48-50). These studies also established that reliable clade designations for *TPA* could be made using an 83-bp region of *tp0548* (48-50), a locus used in epidemiological studies as part of the enhanced *TPA* typing scheme (51, 52) (Fig. S1A). At the outset, *TPA* in clinical samples from all three study sites were amplified using *tp0548*-specific primers; partial sequences were obtained from 29 samples (Fig. S1B). Among the 14 samples from Cali, seven contained *tp0548* type f, placing these strains in the SS14 clade, while the remaining seven contained an assortment of types belonging to the Nichols clade. The six SF strains were a mixture of types within the SS14 clade. Types for seven CZ strains also fell within the SS14 clade, while two contained *TPA* belonged to the Nichols clade.

Nucleotide polymorphisms within *tp0558* (encoding a NiCoT family nickel-cobalt inner membrane permease (53)), also can be used for clade discrimination (48, 54). Based on *tp0558* sequences, clade assignments for *TPA* within 30 clinical samples were determined (Fig. 2). While the majority were exact matches for either clade at all five *tp0558* “discriminator” nucleotide positions, Cali_84, Cali_123, and Cali_133 contained substitutions not found in any of the *TPA* reference strains.

**Figure 1.**
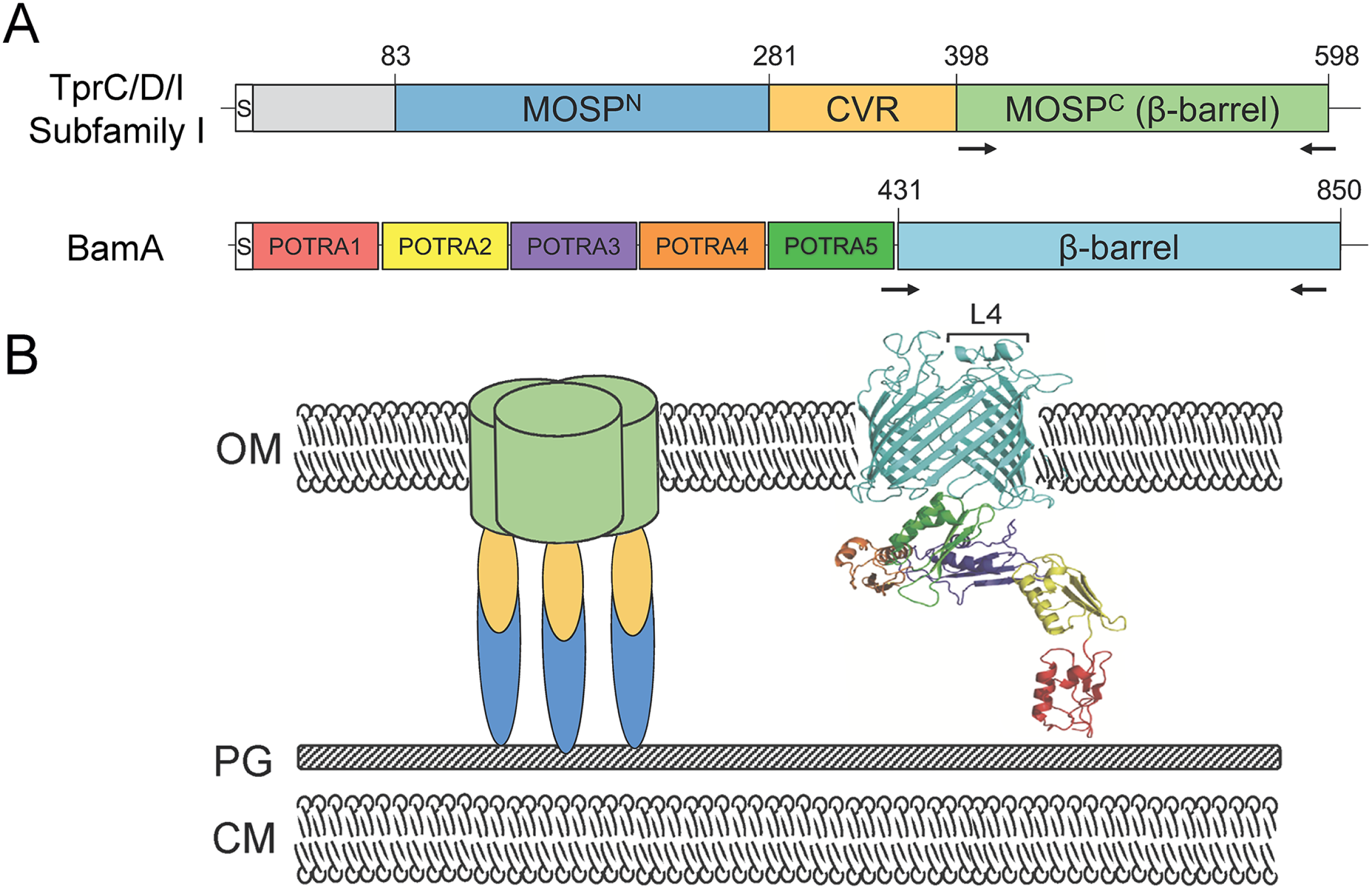
Domain architectures and membrane topologies of TprC (TP0117), TprD (TP0131), TprI (TP0620), and BamA (TP0326). **A**. *T. pallidum* repeat (Tpr) subfamily I paralogs TprC (TP0117), TprD (TP0131), and TprI (TP0620) have identical domain architectures (35, 36). MOSP^N^ and MOSP^C^ correspond to conserved domains shared with the N- and C-termini of the major outer sheath protein (MOSP) of *T. denticola*, the parental Tpr ortholog, identified by the NCBI conserved domain database (CDD) server. Arrows indicate the regions PCR-amplified for sequencing (see Table S3 for primers). Central variable region (CVR) denotes a sequence variable stretch present in all Tprs (40, 77). BamA consists of a C-terminal β-barrel and five periplasmic polypeptide transport-associated (POTRA) domains (34, 37). Numbers refer to amino acid positions within the full-length proteins (signal peptides, denoted by “S”, included) from *TPA* Nichols. **B**. Membrane topologies. In Tprs and MOSP, the MOSP^C^ domain forms the surface-exposed β-barrel (35, 36, 42). Immunofluorescence experiments in *T. pallidum* have confirmed the periplasmic location of MOSP^N^ and the CVR of TprC/D (Nichols) and TprI (35, 36). Moreover, the periplasmic portions of TprC/D and TprI form extended structures, as determined by small angle X-ray scattering analysis, that anchor the β-barrels to the peptidoglycan sacculus (35, 36). A homology model based on the solved structure of the *Neisseria gonorrhoeae* ortholog (38) predicts that the β-barrel of *TPA* BamA contains 16 transmembrane β-strands and eight extracellular loops (37). OM, outer membrane; PG, peptidoglycan; CM, cytoplasmic membrane; L4, BamA immunodominant extracellular loop 4.

**Figure 2.**
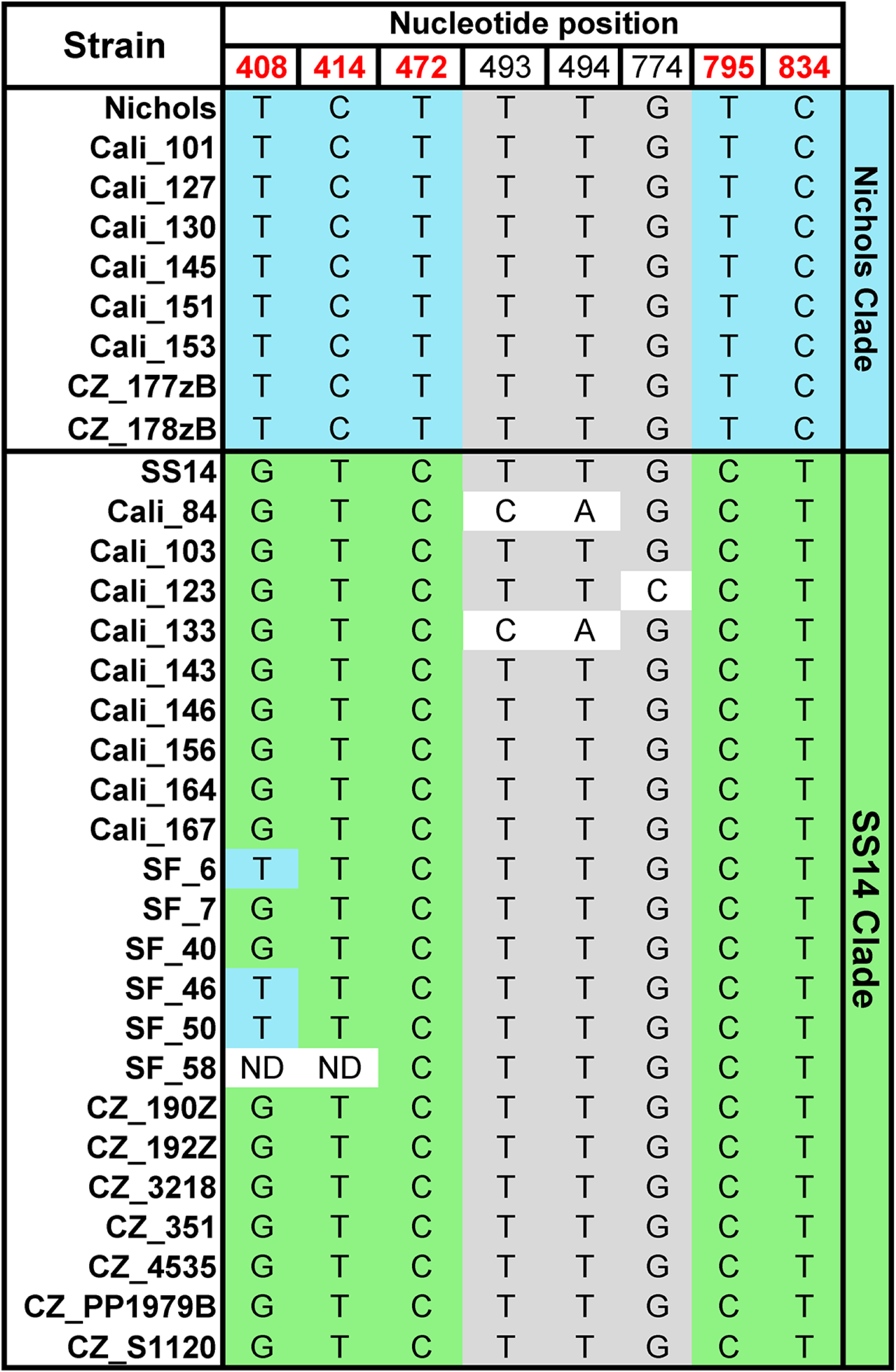
Clade assignments of *TPA* in clinical samples based upon *tp0558* sequences. Nucleotide polymorphisms in Nichols and SS14 *TPA* reference and clinical strains. The five ‘discriminator’ nucleotide positions used for clade assignment (48) are shown in red. Blue and green shading indicate discriminator nucleotides conserved in Nichols and SS14 reference strains, respectively. Grey shading indicates nucleotides conserved in both clades. “ND” indicates nucleotide positions that could not be determined by sequencing of PCR amplicons. All mutations are synonymous except for those at nucleotide positions 493 and 494, which result in the substitution of histidine for phenylalanine (Phe165) in Cali_84 and Cali_133.

In summary, based upon *tp0548* and/or *tp0558*, of the 31 clinical samples, *TPA* within eight clinical samples belonged to the Nichols clade, 21 belonged to the SS14 clade, and two were unassignable because of sequence discordance (Cali_133) or incomplete sequence data (Cali_77) (Table 1).

**Table 1.** TprC, TprD/D2, and BamA allele β-barrel variants and enhanced CDC typing of *TPA* strains in patient specimens from Cali, Colombia, San Francisco, and Czech Republic.

| Clade<sup>1</sup> | Patient | β-barrel variant<sup>2</sup> | | | ECDCT<sup>3</sup> |
| --- | --- | --- | --- | --- | --- |
|  |  | TprC | TprD/D2 | BamA |  |
| Nichols | Cali_101 | Mexico A/PT_SIF | D2 | Mexico A | 14d/d |
|  | Cali_127 | Mexico A/PT_SIF | D2 | Mexico A | 21a/d |
|  | Cali_130 | Nichols | D | ND | XX/d |
|  | Cali_145 | Nichols | D | Nichols (partial) | XX/a |
|  | Cali_151 | Nichols | D | ND | XX/a |
|  | Cali_153 | Sea81-4 | D2 | ND | XX/d |
|  | CZ_177zB | Sea81-4 | D2 | Sea81-4 | 14X/d |
|  | CZ_178zB | Sea81-4 | D2 | Sea81-4 | 14d/d |
| SS14 | Cali_84<sup>4</sup> | Mexico A/PT_SIF | D2 | Nichols | 14d/X |
|  | Cali_103 | Mexico A/PT_SIF | D2 | Mexico A | 12d/f |
|  | Cali_123 | Mexico A/PT_SIF | D2 | Mexico A | 14d/f |
|  | Cali_143 | Mexico A/PT_SIF | D2 | Mexico A | 14d/f |
|  | Cali_146 | Mexico A/PT_SIF | D2 | Mexico A | 14b/f |
|  | Cali_156 | Mexico A/PT_SIF | D2 | SS14 (partial) | XX/f |
|  | Cali_164 | Mexico A/PT_SIF | D2 | Nichols | XX/f |
|  | Cali_167 | Mexico A/PT_SIF | D2 | ND | XX/f |
|  | CZ_190Z | Mexico A/PT_SIF | D2 | SS14 | 14d/k |
|  | CZ_192Z | Mexico A/PT_SIF | D2 | SS14 | 14d/g |
|  | CZ_351 | Mexico A/PT_SIF | D2 | SS14 | 14d/g |
|  | CZ_3218 | Mexico A/PT_SIF | D2 | SS14 | 14e/g |
|  | CZ_4535 | Mexico A/PT_SIF | D2 | SS14 | 14d/g |
|  | CZ_PP1979B | Mexico A/PT_SIF | D2 | SS14 | 14d/g |
|  | CZ_S1120 | Mexico A/PT_SIF | D2 | SS14 | 14d/g |
|  | SF_6 | Mexico A/PT_SIF | D2 | Mexico A | 14d/f |
|  | SF_7 | Mexico A/PT_SIF | D2 | Mexico A | 14d/g |
|  | SF_40 | Mexico A/PT_SIF | D2 | Mexico A | 15a/e |
|  | SF_46 | Mexico A/PT_SIF | D2 | Mexico A | 14d/g |
|  | SF_50 | Mexico A/PT_SIF | D2 | Mexico A | 14d/f |
|  | SF_58 | Mexico A/PT_SIF | D2 | Mexico A | 14d/f |
| Nichols/SS14<sup>5</sup> | Cali_133 | Mexico A/PT_SIF | D2 | Nichols | 14d/y |
| ND<sup>6</sup> | Cali_77 | Mexico A/PT_SIF | ND | Mexico A | 16d/X |

<sup>1</sup>Clade designations based on *tp0548* genotypes (Fig. S1A) and *tp0558* sequences (Fig. 2). See Methods and Table S2 for additional details. The *tp0548* genotype is indicated by the last letter of the ECDCT designation.
<sup>2</sup>The β-barrel-encoding domains of *tprC* from Mexico A and PT\_SIF are identical. Nucleotide alignments for *tprC*, *tprD/D2* and *bamA* β-barrels are presented in Fig. S2, S4 and S5, respectively. See Table S4 for NCBI accession numbers.
<sup>3</sup>Enhanced CDC typing, performed as described in Materials and Methods. Strain type designations are as follows: the first numeral represents the number of repeats in the *arp* gene; the first letter represents the *MseI* restriction site profile in the *tprE/G/J* genes; and the second letter is based on sequence analysis of an 83-bp region of *tp0548*. X's are used to indicated missing data for a given position.
<sup>4</sup>Clade designation based on *tp0558* alone.
<sup>5</sup>Encodes a unique Nichols-like *tp0548* (genotype y) and an SS14 *tp0558*. See Fig. S1A for additional details.
<sup>6</sup>ND, not determined due to limited sample material.

### Classification of TprC β-barrel alleles encoded by *TPA* reference strains

Nucleotide sequence comparisons of full-length *tprC* (*tp0117*) genes encoded by the 41 *TPA* reference genomes available, examined using fastx_collapser, identified five alleles (Table S2). Four of these are represented by reference strains Nichols, SS14, Mexico A and Seattle81-4 (Sea81-4) (40, 48, 55), while the fifth is represented by the cohort of 25 clinical strains (designated by PT_SIF) from Lisbon, Portugal (56). The five *tprC* alleles encode four β-barrel variants (Fig. 3A and B); the Mexico A and PT_SIF β-barrel-forming domains are identical (Fig. S2) and, therefore, are together referred to as Mexico A/PT_SIF. The nucleotide polymorphisms that distinguish the four TprC β-barrel variants are distributed over four regions (Fig. 3A and B; Fig. S2). Region I consists of a 56-nucleotide stretch in which seven positions are unique to the Nichols allele. Region II contains the sole nucleotide substitution that differentiates the Mexico A/PT_SIF and SS14 β-barrel alleles. Region III consists of a 19-nucleotide stretch in which seven positions are unique to the Sea81-4 β-barrel allele. Region IV consists of a 49-nucleotide stretch in which seven positions are shared by Nichols and Sea81-4. Thirteen of the 22 polymorphisms dispersed across the β-barrel domain result in amino acid substitutions, six of which are non-conservative (Fig. 3B). The ‘branch-site’ model in the phylogenetic analysis by maximum likelihood (PAML) package (57) identified six amino residues in full length *tprC* as being positively selected (P > 95%), all of which were in the β-barrel domain (Fig. 3A).

**Figure 3.**
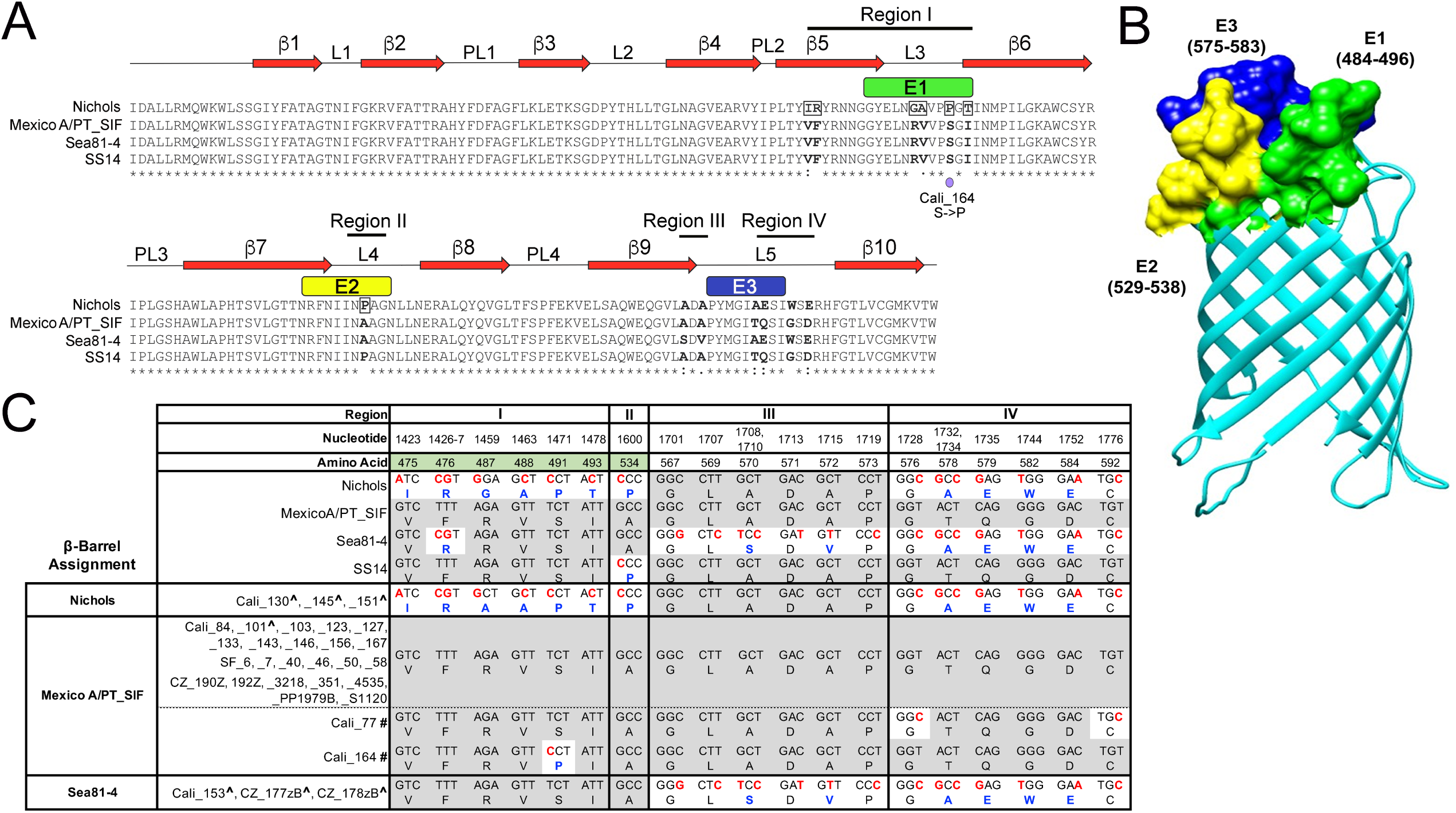
*TPA* in clinical samples from Cali, San Francisco, and the Czech Republic encode Nichols, Mexico A/PT_SIF or Sea81-4 TprC allele β-barrel domains. **A.** Multiple sequence alignment of the four TprC allele β-barrels identified in *TPA* reference genomes. Identical amino acids are indicated by asterisks (*); non-synonymous amino acid substitutions are in bold. Double (:) and single dots (.) indicate highly conservative and conservative changes. Positively selected amino acids by branch-site model using PAML (57) are boxed in the Nichols allele sequence. Numbered red arrows and black lines indicate the positions of predicted β-strands, periplasmic loops (PL), and extracellular loops (L) in the Nichols TprC allele β-barrel structural model shown in panel C. A purple dot indicates a non-synonymous nucleotide substitution in the Mexico A allele β-barrel variant encoding region for Cali_164. **B**. The TMBpro webserver (58) was used to generate a three-dimensional structural model for the β-barrel of TprC (Nichols). In all three panels, B-cell epitopes E1 (residues 484-496), E2 (residues 529-538) and E3 (residues 575-583) predicted by DiscoTope 2.0 (59) are shown in green, yellow, and dark blue. **C**. Nucleotide and amino acid differences, shown in red and blue, respectively, in the TprC allele β-barrels and variants identified in *TPA* reference genomes and 31 clinical strains, respectively. The Mexico A and PT_SIF TprC allele β-barrel sequences are identical and referred to as Mexico A/PT_SIF. The nucleotide and amino acid numbers for variable positions are based on full-length *TPA* Nichols TprC (Fig. S2). Consensus positions are shaded gray. Positively-selected amino acids identified by branch-site model using PAML (57) are shaded green. Carat symbols (^) indicate clinical strains belonging to the Nichols clade (see Table 1). Hash marks (#) designate Mexico A allele TprC β-barrel variants (Cali_77 and Cali_164).

### Structural modeling of the TprC β-barrel and topological mapping of the amino acid substitutions that differentiate the four β-barrel variants

To date, there are no solved structures for any of the Tpr proteins. As a first step towards understanding the topological and/or functional implications of the above sequence data, a structural model for the Nichols TprC β-barrel was generated using TMBpro (58). As shown in Fig. 3B, the predicted β-barrel consists of ten anti-parallel β-strands with five connecting extracellular loops of varying size. Overall, this model is consistent with the general principles of amphipathic β–barrel structure (10, 13) in that the external surface facing the lipid bilayer is highly hydrophobic, while charged residues line the channel (Fig. 4). It is interesting to note that the strong positive charge within the channel could explain its high conductivity for the fluorophore Tb(DPA)_3_^3-^ used in previous studies of MOSP^C^ domain porin activity (35, 36). Also consistent with general OMP structure are the relatively large extracellular loops and short periplasmic turns. Depending on location, substitutions in the barrel could affect either the properties of the aqueous channel or surface interactions between the treponeme and its obligate human host. Nine of the 13 amino acid polymorphisms in the four TprC β-barrel domain variants are in extracellular loops (three in L3, one in L4 and five in L5), while four are in predicted transmembrane strands (including two positively predicted residues in β5) (Fig. 3A).

**Figure 4.**
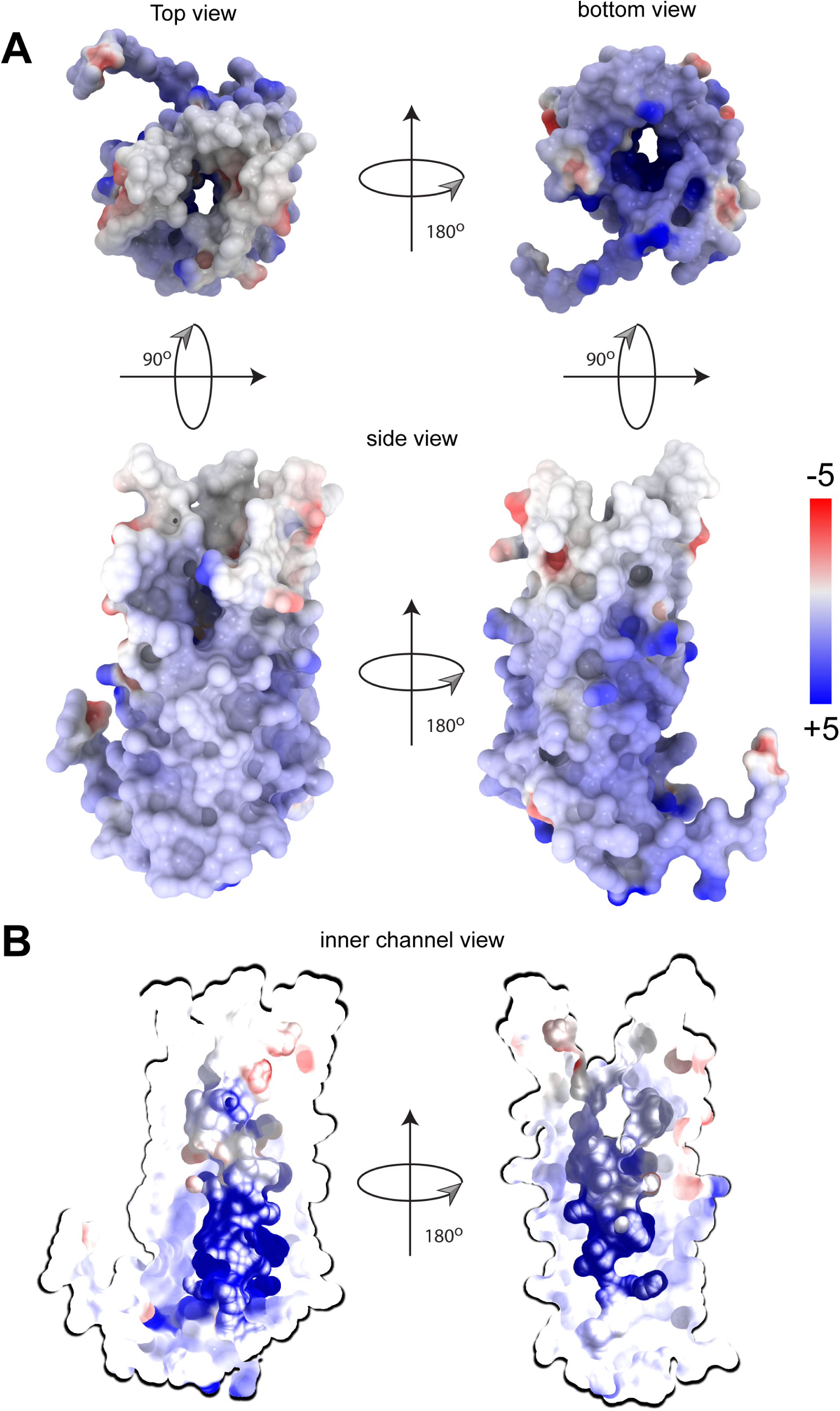
Nichols TprC/D allele β-barrel channel interior is highly positively-charged. **A**. Structural model of the Nichols TrpC/D allele β-barrel shown as a surface colored by electrostatic potential. Positively and negatively charged residues are shown in blue and red, respectively. **B**. A slice through the Nichols TrpC/D allele β-barrel porin channel reveals a charged interior pore. The pore is colored according to its charge distribution scale shown on the right. The electrostatic potential display was generated using ICM MolBrowserPro (97).

### The majority of B-cell epitopes predicted for TprC localize to extracellular loops

Immunodominant B-cell epitopes of Gram-negative OMPs typically are located in extracellular loops and often are sequence-variable (14-18). Analysis of the TprC β-barrel structural model using DiscoTope 2.0 (59) revealed that the predicted B-cell epitopes align well with extracellular loops L3, L4, and L5 (Fig. 3B), and closely correspond to variable regions I, II, and III/IV, respectively (Fig. 3A). Importantly, each predicted B-cell epitope contains at least one non-conservative substitution (Fig. 3A). Of note, the single amino acid residue (Pro534) that differentiates the Mexico A/PT_SIF and SS14 alleles (Region II) occurs in predicted epitope E2 (Fig. 3A).

### *TPA* strains within clinical samples from Cali, San Francisco, and the Czech Republic contain three TprC β-barrel variants, with Mexico A/PT_SIF predominating

TprC β-barrel sequences from all 31 clinical samples were obtained (Table 1 and Fig. 3C). Overall, the Mexico A/PT_SIF β-barrel variant predominated (25/31). While Cali_77 and Cali_164 contained nucleotide changes in Regions IV and I, respectively, they were designated as Mexico A/PT_SIF-like because they contain the ‘differentiator’ G at nucleotide position 1600 (Region II). Cali_164 contained a proline in place of a serine at residue 491 in Region I (L3/E1; Fig. 3C), thereby creating a novel E1 epitope. The remaining TprC β-barrel variants within the clinical samples were exact matches for either the Nichols or Sea81-4 allele. Notably, of the eight *TPA* clinical strains assigned to the Nichols clade (indicated by carat symbols in Fig. 3C), only three contained a Nichols β-barrel variant, while all of the clinical strains assigned to the SS14 clade contained a Mexico A/PT_SIF β-barrel variant (Table 1 and Fig. 3C). Cali_133, the strain with discordant *tp0548* and *tp0558* sequences, also contained a Mexico A/PT_SF β-barrel variant.

### *TPA* reference strains encode only two TprD β-barrel variants

Centurion-Lara and co-workers (40, 60) were the first to report that the *tp0131* locus can harbor either a *tprD* (which is identical to *tprC* in the Nichols reference strain) or *tprD2* allele. As noted earlier (40), the *tprD* allele occurs only in reference genomes with a Nichols *tprC* allele (Table S2). Alignment of full-length *tprD* and *tprD2* from the reference strains (Fig. S3) reveals much greater sequence divergence than *tprC* (Fig. S2); *tprD* and *tprD2* encode identical MOSP^N^ domains but divergent central variable regions (CVR) and MOSP^C^ domains (Fig. S3). Comparison of the TprD and TprD2 β-barrel-encoding (MOSP^C^) domains revealed four regions of variability (Fig. 5A and S3). Region I consists of a single nucleotide change that results in substitution of arginine for lysine. Regions II-IV contain numerous nucleotide polymorphisms, many of which result in non-conservative amino acid substitutions. Region III also contains two single nucleotide in-dels.

**Figure 5.**
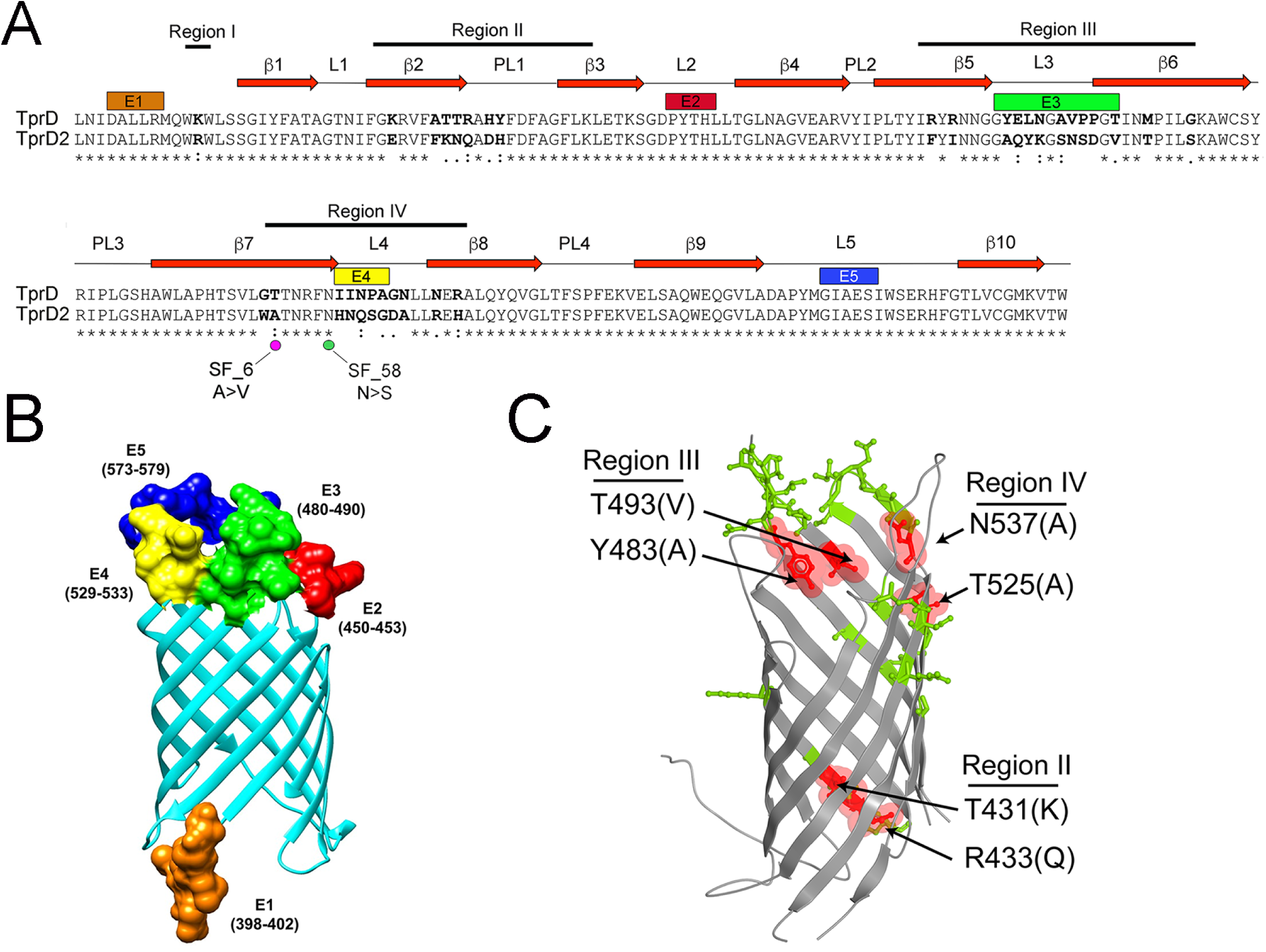
*TPA* in clinical samples from Cali, San Francisco, and the Czech Republic encode TprD or D2 allele β-barrels. **A.** Pairwise alignment of TprD and D2 allele β-barrels identified in *TPA* reference genomes. Identical amino acids are indicated by asterisks (*). Non-synonymous amino acid substitutions are in bold. Double (:) and single dots (.) indicate highly conservative and conservative changes. Numbered red arrows and black lines indicate the positions of predicted β-strands, periplasmic loops (PL) and extracellular loops (L) 1-5 in the TprD2 structural model shown in panel B. Magenta and green dots, respectively, indicate non-synonymous nucleotide changes and amino acid substitutions in clinical strains SF_6 and SF_58. **B**. The TMBpro web server (58) was used to generate a three-dimensional structural model for the TprD2 allele β-barrel. In panels A and B, B-cell epitopes E1 (residues 398-402), E2 (residues 450-453), E3 (residues 480-490), E4 (residues 529-533) and E5 (residues 573-579) predicted by DiscoTope 2.0 are shown in orange, red, green, yellow, and dark blue, respectively. **C**. Ribbon diagram comparing the TprD and TprD2 allele β-barrels. Residues that differ between TprD and TprD2 are shown as sticks. Variable residues predicted to affect the pore opening and exit are highlighted in red and labeled to indicate the corresponding variable Region. Conserved residues are shown in gray. Residue numbers correspond to the full-length Nichols TprD.

### Topological mapping of predicted B-cell epitopes and sequence differences used to distinguish TprD and D2 allele β-barrels

Based on the predicted structural model (Fig. 5B), Region I lies in the extreme N-terminus of the β-barrel. Region II encompasses β-strands β2 and β3 and the intervening periplasmic loop PL1, while Regions III and IV center about predicted extracellular loops L3 and L4, respectively. Of the five B-cell epitopes predicted by DiscoTope 2.0, four reside in extracellular loops (Fig. 5A and B). Of note, L2, which coincides with epitope E2, is conserved between TprC, TprD, and TprD2 (Fig. 3A and 5A). Whereas L5 in TprC is variable (Fig. 3A), this loop and corresponding epitopes (E3 and E5, respectively) are identical in TprD and D2 (Fig. 5A). In addition to amino acid differences in the extracellular loops, the models for TprC/D and TprD2 identified amino acid substitutions at the entrance and exit of the channel (Fig. 5C, highlighted in red) with the potential to affect porin functionality.

### TprD2 allele β-barrels predominate in the clinical strains

*TPA* strains encoding TprD2 allele β-barrel predominated in the 30 clinical samples from which *tprD* sequences were obtained (Table 1). Of the eight strains assigned to the Nichols clade, only Cali_130, Cali_145, and Cali_151 contained Nichols allele. The remaining five strains belonging to the Nichols clade, all 21 belonging to the SS14 clade, and one indeterminate (Cali_133) contained a TprD2 β-barrel variant. The β-barrel-encoding sequences of SF_6, and SF_58 harbored single nonsynonomous nucleotide substitutions in their predicted β7 strands (Fig. 5A); given the imprecision of β-strand prediction, it is possible that the nonconservative change in SF_58 occurs in extracellular loop L4 (epitope E4).

### Classification of BamA alleles encoded by *TPA* reference strains

Analysis of full-length *bamA* genes from all available *TPA* reference genomes using fastx_collapser identified four alleles, represented by Nichols, SS14, Mexico A, and Sea81-4 (Table S2 and Fig. S4). The sequence differences that distinguish these alleles are restricted to six variable regions within the β-barrel (Fig. S4) and result in 13 amino acid changes (seven non-conservative) and a five amino acid deletion in the Mexico A allele (Fig. 6A and B). Region I contains a 49-nucleotide stretch unique to the SS14 allele. Region II consists of a single nucleotide change found only in the Mexico A allele. Region III also consists of a single nucleotide with a ‘G’ in the Mexico A and SS14 alleles and a ‘T’ in Nichols and Sea81-4 alleles. Region IV, the most polymorphic, consists of either a single nucleotide substitution along with a 15-nucleotide deletion unique to the Mexico A allele or a non-identical stretch of 15 nucleotides in the other three alleles; of note, substitutions in this region modify the polyserine tract (Fig. 6A and B; Fig. S4), a unique feature of BamA orthologs in pathogenic treponemes (34, 61, 62). Regions V and VI consist of single nucleotide changes specific to the Mexico A and SS14 alleles, respectively. The ‘branch-site’ model (57) did not identify any positively selected amino acids in the β-barrel region; however, the ‘site’ model identified six positively selected amino acids (P > 95%), distributed over Regions I-IV (Fig. 6A and B).

**Figure 6.**
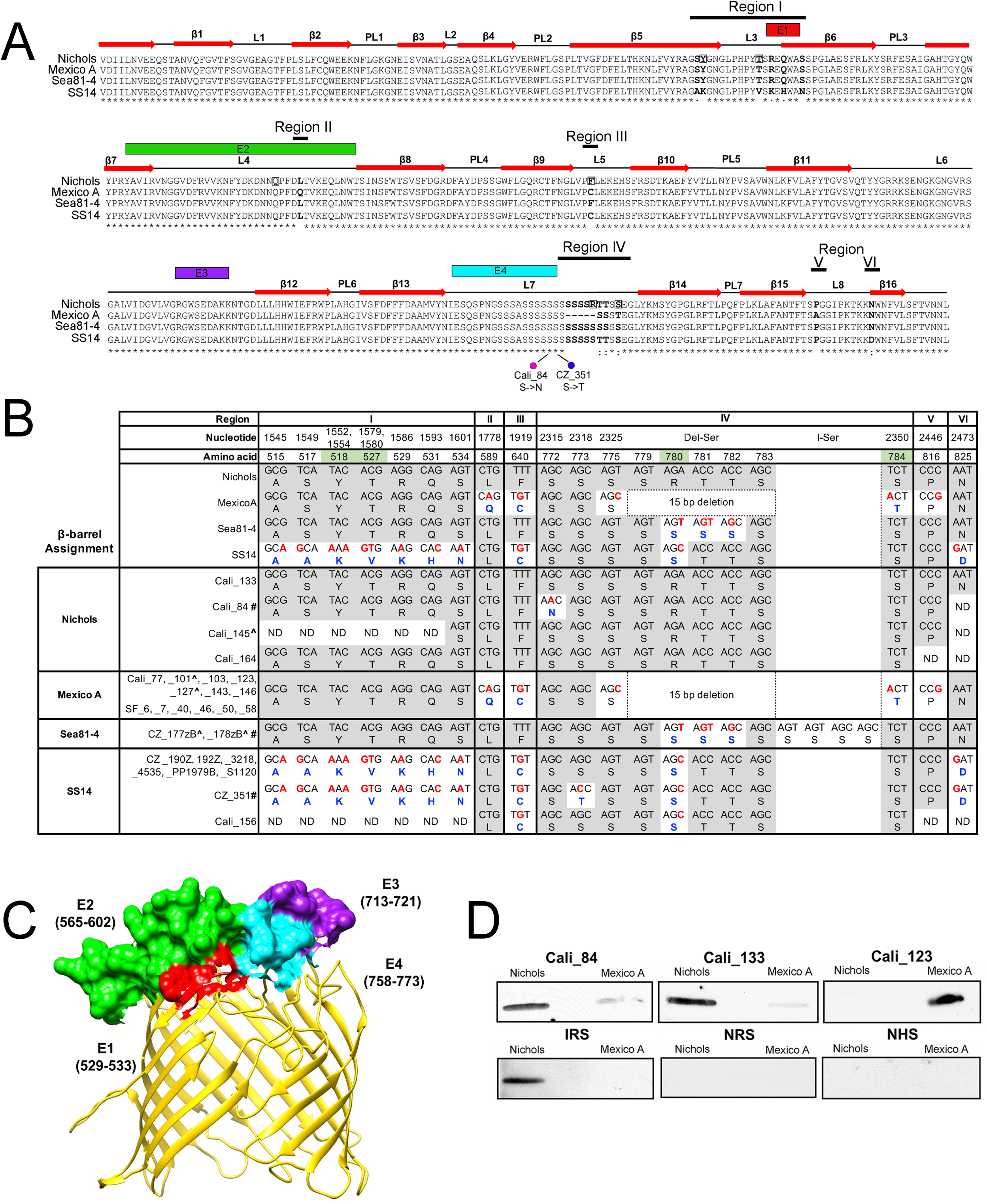
*TPA* in clinical samples from Cali, San Francisco, and the Czech Republic encode Nichols, Mexico A, Sea81-4 and SS14 BamA allele β-barrel variants. **A.** Multiple sequence alignment of the four BamA allele β-barrels identified in *TPA* reference genomes. Identical amino acids are indicated by asterisks (*); non-synonymous amino acid substitutions are in bold. Double (:) and single dots (.) indicate highly conservative and conservative changes, respectively. Positively selected amino acids by the site model using PAML (57) are boxed. Numbered red arrows and black lines indicate the positions of β-strands, periplasmic loops (PL), and extracellular loops (L) in the Nichols BamA structural homology model (37) shown in panel B. Magenta and green circles indicate nonsynomous substitutions in the Cali_84 (Nichols) and CZ_351 (SS14) BamA β-barrels, respectively. B-cell epitopes E1 (residues 529-533), E2 (residues 565-602), E3 (residues 713-721), and E4 (residues 758-773), predicted by DiscoTope 2.0, are shown in red, green, purple, and cyan boxes, respectively. **B.** Nucleotide and amino acid differences, shown in red and blue, respectively, in the BamA allele β-barrels and variants found in *TPA* reference genomes and 27 clinical strains. Nucleotide and amino acid numbers for variable positions are based on full-length Nichols BamA (Fig. S4). The column designated ‘Del-Ser’ indicates a 15-bp deletion within the Mexico A BamA allele β-barrel. The column designated ‘I-Ser’ indicates a 12-nucleotide insertion in CZ_177zB and CZ_178zB that extends the polyserine tract by 4 residues. Consensus positions are shaded gray. Positively selected amino acids identified by branch-site model are shaded green. ‘ND’ indicates nucleotide and amino acid positions that could not be determined by sequencing of PCR amplicon. Carat symbol (^) indicates clinical strains assigned to the Nichols clade (see Table 1). Hash marks (#) designate BamA β-barrel variants encoded by Cali_84 (Nichols), Cz177zB (Sea81-4), Cz178zB (Sea81-4) and CZ_351 (SS14). **C.** Structural homology model for the Nichols BamA allele β-barrel showing the position of B-cell epitopes using the same color scheme as in panel A. **D.** A non-conservative amino acid substitution in L4 of BamA markedly alters the reactivity of syphilis patient sera. Sera from Cali_84 and Cali_133, patients infected with strains containing Nichols BamA β-barrel variants, preferentially recognize Nichols L4 while serum from Cali_123, infected with a strain containing a Mexico A allele BamA β-barrel, preferentially recognizes Mexico A L4. Each lane contains ~100 ng of affinity-purified recombinant histidine-tagged protein. NRS, normal rabbit serum; IRS, immune rabbit serum obtained from a rabbit infected with *TPA* Nichols; NHS, normal human serum.

### Topological mapping of the amino acid substitutions used to distinguish between BamA alleles β-barrels

We previously described and partially validated by immunofluorescence analysis a structural homology model for *TPA* BamA consisting of 16 transmembrane β-strands and eight extracellular loops (37). Variable regions I through V are located entirely or largely in extracellular loops (Fig. 6A). The DiscoTope 2.0 server predicts that extracellular loops L4, L6 and L7 contain major epitopes (E2, E3, and E4, respectively; Fig. 6A and C). It is interesting to note that variable region IV in L7 lies outside of E4, raising the possibility that immune pressure may not be driving variability in this loop.

### The clinical strains encode all four BamA β-barrel variants

β-barrel sequences matching all four of the BamA reference alleles were identified in 27 clinical samples (Table 1 and Fig. 6B). However, several varied from their corresponding reference (Fig. 6B). Most notably, the polyserine tract of CZ_177zB and CZ_178zB, both of which encoded Sea81-4-like BamA allele β-barrels, contained a 12-nucleotide insertion that adds four additional serine residues (Fig. 6B). *bamA* β-barrel sequences were obtained from twenty-five *TPA* clinical strains with clade designations (Table 1). Of the five assigned to the Nichols clade, only Cali_145 contained a Nichols BamA β-barrel variant, while Cali_101 and Cali_127 contained Mexico A β-barrel variants. Of the 20 strains belonging to SS14 clade, only eight contained an SS14 β-barrel variant; of the remaining 12, ten encoded the Mexico A *bamA* β-barrel variant and two had Nichols variants.

### A single amino acid substitution in BamA extracellular loop 4 alters immunoreactivity by patient sera

As noted above, the nucleotide change in variable Region II results in an amino acid substitution (glutamine for leucine at residue 589) unique to the Mexico A allele β-barrel. We recently reported that this polymorphism markedly alters the reactivity of patient sera with a heterologous L4 peptide (37). Immunoblots performed with sera from Cali patients infected with *TPA* strains containing Nichols (Cali_84 and Cali_133) or Mexico A BamA β-barrel variants (Cali_123) confirmed this finding (Fig. 6D).

### β-barrel profiles in clinical strains can be mosaics of reference genome profiles

Clade distributions and “across the board” β-barrel profiles of clinical strains were compared with those of the *TPA* reference genomes (Table S2). Four of the six reference strains belong to the Nichols clade (CDC A, Nichols, Chicago, and DAL-1) had Nichols *tprC*, *tprD,* and *bamA* alleles, while Sea81-4 and UW189B harbored *tprD2* in addition to Sea81-4 *tprC* and *bamA*. Although all members of the SS14 clade harbored *tprD2* alleles, variability at the other two loci, particularly *tprC,* was noted (Table S2). Parsimony analysis (63) of the *TPA* reference genomes confirmed that the three OMP loci were poor predictors of clade assignment. *tprC, tprD,* and *bamA* contained only 25.9%, 0%, and 5.6% parsimony informative sites, respectively, compared to 80.6% for *tp0548* and 100% for *tp0558.*

Sequences from all three OMP loci in five clinical strains assigned to the Nichols clade were obtained (Tables 1 and 2). Of these, only Cali_145 possessed an OMP profile resembling that of the Nichols reference strain. The profiles of CZ_177zB and CZ_178zB were Sea81-4-like, while Cali_101 and Cali_127 had Mexico A-like profiles. Thus, two of five Nichols clade strains had β-barrel variant profiles matching *TPA* belonging to the SS14 clade. Complete profiles were obtained for 20 strains assigned to the SS14 clade (Tables 1 and 2). Strikingly, although all had TprD2 β-barrel variants, none had profiles matching the SS14 reference strain. All had the Mexico A/PT_SIF TprC β-barrel variant, while only eight had the SS14 strain BamA allele β-barrel variant. The other BamA β-barrel variants were either Mexico A (n = 10) or Nichols (n = 2). When stratified by geographic location (Table 1), the uniformity of the SF strains contrasted with the diversity of strains from Cali.

**Table 2.** Summary of β-barrel allele profiles and molecular types in clinical samples with complete OMP profiles and belonging to either the Nichols or SS14 clade.

| Clade | Number of samples | *tprC* β-barrel | *tprD/D2* β-barrel | *bamA* β-barrel | Reference Genome OMP profile<sup>1</sup> | ECDCT<sup>2</sup> |
| --- | --- | --- | --- | --- | --- | --- |
| Nichols | 1 | Nichols | D | Nichols | Nichols | XX<sup>3</sup>/a (1) |
|  | 2 | Mexico A/<br>PT_SIF | D2 | Mexico A | Mexico A | 14d/d (1)<br>21a/d (1) |
|  | 2 | Sea81-4 | D2 | Sea81-4 | Sea81-4 | 14d/d (1)<br>14X/d (1) |
| SS14 | 10 | Mexico A/<br>PT_SIF | D2 | Mexico A | Mexico A | 12d/f (1)<br>14d/f (5)<br>14b/f (1)<br>14d/g (2)<br>15a/e (1) |
|  | 8 | Mexico A/<br>PT_SIF | D2 | SS14 | Amoy | XX/f (1)<br>14d/k (1)<br>14d/g (5)<br>14e/g (1) |
|  | 2 | Mexico A/<br>PT_SIF | D2 | Nichols | Unique | 14d/X (1)<br>XX/f (1) |

<sup>1</sup>Based on matching the reference genome at all three OMP loci.
<sup>2</sup>Number in parenthesis indicates that number of clinical samples containing *TPA* of the corresponding ECDC type.
<sup>3</sup>X, Undetermined genotype

### Molecular typing does not correlate with β-barrel profiles

The enhanced CDC typing (ECDCT) system has been widely used to study the diversity and epidemiology of *TPA* strains in numerous global locales (64, 65). To determine whether molecular typing is predictive of OMP profiles, ECDCT was completed on *TPA* strains in 21 clinical samples (Tables 1 and 2). Altogether, 10 different ECDCTs were detected. 14d/g (n = 7) and 14d/f (n = 5) were the most prevalent and, along with 14d/d (n = 2), the only genotypes found at more than one site. Five strains had an ECDCT (14d/f) matching the SS14 reference strain but Mexico A-like OMP profiles; of the 12 strains with Mexico A allele β-barrels in all three OMP loci, none matched the Mexico A reference strain genotype (16d/e). Two ECDCTs (14d/d and 14d/g) were associated with more than one OMP profile. Conversely, several OMP profiles were associated with more than one ECDCT. Collectively, these indicate that OMP profiles of *TPA* strains cannot be predicted based on ECDCT data.

## DISCUSSION

For decades, syphilologists have sought means to distinguish strains of *TPA* for epidemiologic, pathogenesis- and vaccine-related investigations. Serologic analyses of live “street strains” by Turner and Hollander in the 1950s (55) yielded evidence for antigenic differences, presumably attributable to surface-exposed epitopes. The molecular typing method introduced by Pillay *et al.* in 1998 (52), based on number of repeats in the *arp* (*tp0433*) gene and sequence polymorphisms in the Tpr subfamily II genes *tprE* (*tpr0313*), *tprG* (*tp0317*) and *tprJ* (*tp0621*), was a major advance, although subsequent studies revealed that this system insufficiently distinguishes common *TPA* strains circulating globally (51, 64). The addition of subtyping based on sequences from *tp0548* markedly improved the discriminatory power of the CDC typing scheme (51, 65). In parallel, Šmajs and co-workers (48, 49) found that *tp0548* is one of several loci that phylogenetically separate *TPA* strains into two clusters, named for the Nichols and SS14 reference strains, and that *tp0548* subtypes segregate into these two clades. Most recently, phylogenetic analysis of whole genome sequences of geographically diverse *TPA* strains has lent strong support to the concept of two clades, with strains belonging to the SS14 clade supplanting those belonging to the Nichols clade (50). Consistent with this notion, the majority of *TPA* in clinical samples examined in this study grouped with the SS14 clade. They also revealed, however, that *TPA* within the Nichols clade are still actively circulating in the Eastern and Western hemispheres.

Arora *et al*. (50) reported that the SS14 clade contains a central dominant haplotype, designated SS14-Ω, which does not include the Mexico A strain (also classified within the SS14 clade). The observation that SS14 clade members in both Cali and San Francisco have a preponderance of Mexico A allele β-barrel at all three OMP loci suggests that Mexico A-like strains are circulating more widely than their analysis suggests. An alternative possibility, which sequence data do not rule out, is that the SS14 clade strains in the study cohort fall within SS14-Ω but some contain Mexico A-like OMP allelic remnants. Exclusion of *bamA* sequences, along with the absence of *tpr* genes in draft genomes used by Arora *et al.* (50), precludes a direct comparison of their data with those presented here.

Phylogenetic reconstructions, including those used to distinguish *TPA* clades (48, 50), rely upon genes whose sequence variation recapitulates the vertical evolution of the bacterium (66). OMP-encoding genes typically are excluded from such analyses because they undergo mutations and rearrangements that give rise to phylogenetic trees in conflict with those derived from “reference” genes (66). However, sequence variation of OMPs to enhance environmental fitness, virulence, and immune evasion plays a central role in the evolution and epidemiology of pathogenic bacteria (15-17, 67, 68). Parsimony analysis revealed that the evolutionary histories of *tprC*, *tprD*, and *bamA* diverge greatly from those of genes used for clade differentiation (*i.e*., *tp0548* and *tp0558*). Inspection of the three OMP genes/proteins in reference and clinical strains explains this dichotomy and provides evidence that OMPs in *TPA* are subject to host-driven, adaptive mechanisms. Except for a few nucleotides in the N-terminal periplasmic portion of Mexico A *tprC*, the variable regions for *tprC* and *bamA* reside within the surface-exposed, OM-embedded β-barrel, just as one would expect if selection pressures exerted by the host are driving variation.

Sequence variation within the β-barrel encoding regions of *tprC*, *tprD,* and *bamA* ranged from point mutations with single amino acid substitutions to small stretches of DNA encoding multiple amino acid differences. Gray *et al.* (69) contended that the variable regions in *tprC* containing multiple amino acid changes are more consistent with small ‘site-specific’ gene conversion events than accumulated point mutations, although they were uncertain whether the source(s) of the acquired sequences is intra-or inter-genomic. Regardless, there is no reason why recombinatorial mechanism(s) would be limited to *tprC*. Indeed, the mosaic OMP profiles in some clinical strains can best be explained by recombination/conversion of larger DNA fragments within and between clades. It is worth noting that genomic sequencing identified a number of OMP loci in pathogenic treponemes, including BamA in *TPA*, where recombination appears to have occurred (50, 70-72). The most plausible scenario for inter-genomic exchange during human syphilis would be anogenital ulcers co-infected with *TPA* strains containing genetically divergent OMP loci.

According to structural models, the variable regions in *TPA* OMP β-barrels coincide with extracellular loops predicted to contain B-cell epitopes. Immunoblot results for L4 of BamA underscore the antigenic impact of even single non-synonymous amino acid substitutions in a surface-exposed region. Substitutions and antigenic diversity in the predicted extracellular loops for Tpr β-barrels likewise could serve in a similar role in immune evasion. The cumulative effect of the OMP loop variants noted in this study, along with those in TprK (56, 65, 73) and as yet uncharacterized *TPA* OMPs (33, 40), would be to foster strain diversity of among *TPA* strains and, thus, their ability to persist at the population level. There are numerous examples of virulence-related OMPs in bacterial pathogens playing a role in maintenance of cellular homeostasis and/or outer membrane integrity (19-21, 23, 25, 74-76). One also must consider the possibility, therefore, that extracellular loop variants impact interactions at the host-pathogen interface during syphilitic infection. The hotspot for variation in extracellular loop L7 of BamA, which lies outside the adjacent predicted major epitope (E4), might be one example.

TprD and TprD2 contain identical MOSP^N^ domains but highly divergent CVRs and β-barrels (40, 69, 77). Thus, although involving only two alleles, generation of diversity at the *tprD* locus was a more complex process than for *tprC* and cannot be explained solely by events shaping the β-barrels. Moreover, since the CVR is periplasmic (36), non-immunologic selection pressures also must have been at work. Comparison of *tprD* and *D2* suggests that variation in the β-barrel likely arose from gene conversion events involving relatively small segments of DNA similar to those proposed for *tprC* (69). However, comparison of the variable regions in the *tprC* and *tprD/D2* loci (Fig. S6) brings to light important differences. While separate conversions involving L3 appear to have occurred in the *tprC* and *tprD*, two exchanges occurred only in the *tprD* locus: Region II, spanning transmembrane strands β2 and β3, and Region IV, containing L4 plus flanking DNA from strands β7 and β8. These differences could affect not only the corresponding predicted extracellular loops of TprD2 but also its porin channel function. Differences in substrate preference that broaden or enhance capacity to import water-soluble nutrients across the outer membrane theoretically would be of great benefit to an extreme auxotroph and obligate pathogen such as *TPA* (27, 77). Additional point mutations, such as those in the TprD2 β-barrels of SF_6 and SF_58 might further fine-tune channel functionality. The current worldwide predominance of strains containing *tprD2* could reflect, at least in part, the greater fitness conferred by the presence of functionally and antigenically distinct TprC and TprD OMPs.

Compared to OMPs of many dual membrane pathogens (11, 18, 78-81), including the sexually transmitted organisms *Neisseria gonorrheae* (82, 83) and *Chlamydia trachomatis* (84, 85), a surprisingly small number of variants for each *TPA* OMP were found. The high degree of similarity between the β-barrel sequences of *TPA* in clinical samples and their reference allele counterparts argues that this limited diversity cannot be explained solely by the comparatively small number of *TPA* genomes sequenced to date. Two possible explanations, which are not mutually exclusive, can be envisioned. One is that *TPA* OMPs are subject to ‘uneven’ immune pressure due to microbiologic factors, such as low copy numbers and/or heterogeneous expression within spirochete populations (33, 77), and variability of antibody responses in persons with different genetic backgrounds. A second possibility is that structural constraints counter-balance immunological forces promoting loop diversity. The notion of favored loop sequences that protect the bacterium has been invoked to explain the preservation of sequence types of PorB, the dominant porin of *Neisseria meningitidis*, commonly associated with invasive disease in surveys of endemic and epidemic meningococcal strains (18).

The worldwide resurgence of syphilis has kindled a sense of urgency for vaccine development (2, 4, 86). To be efficacious globally, a syphilis vaccine must target surface-exposed (*i.e.*, antibody-accessible) determinants expressed by geographically disparate *TPA* strains. Data presented herein make clear that vaccine development based on integral OMPs needs to proceed along two broad fronts. One is to expand the list of candidate vaccinogens through topological and structural characterization of proteins known or predicted to form outer membrane-embedded β-barrels (33), coupled with assessment of opsonophagocytosis activity *ex vivo* and protection in the experimental rabbit model (86). The second is to refine methods for genomic sequencing of *TPA* strains in clinical samples (56) to catalog sequence diversity among individual OMPs on a global level. From a vaccine standpoint, the objective would be to develop a mono- or multi-valent vaccine based on OMP alleles from *TPA* strains that are circulating in at risk populations worldwide. Similar vaccine strategies have been proposed for *Borrelia burgdorferi*, the Lyme disease spirochete (87-89). The ability to use conserved extracellular loops conserved (*e.g.*, L2 in TprC/D/D2, L5 in TprD/D2, and L3 in BamA) would circumvent the difficulties associated with expression and purification of full-length OMPs on a mass scale (70).

## MATERIALS AND METHODS

### Clinical Samples

Blood and skin biopsies were obtained during 2009-2014 from patients with untreated secondary syphilis identified and referred for enrollment through a previously described network of health care professionals in Cali, Colombia (43, 44) according to protocols approved at Centro Internacional de Entrenamiento e Investigaciones Médicas (CIDEIM). Swabs from primary syphilis lesions were obtained during 2004-2007 at the San Francisco Municipal STD Clinic according to protocols approved by the University of California, San Francisco and the Centers for Disease Control and Prevention (45). Swabs from primary and secondary syphilis lesions were obtained during 2012-2013 in the Czech Republic at Brno (Department of Dermatovenereology, St. Anne’s Faculty Hospital, Masaryk University) and Prague (National Reference Laboratory for the Diagnostics of Syphilis, The National Institute of Public Health) (48) in accordance with protocols approved by the Ethics Committee of the Faculty of Medicine, Masaryk University. Samples and clinical data were de-identified at the sites of origin prior to transmission to UConn Health.

### Propagation of *TPA* and generation of immune rabbit serum (IRS)

Animal protocols were approved by the UConn Health Institutional Animal Care and Use Committee under the auspices of Animal Welfare Assurance A347-01. *TPA* Nichols was propagated by intratesticular inoculation of New Zealand white rabbits (90). To generate IRS, animals were challenged intradermally with 1 × 10^3^ treponemes at each of six dorsal sites ~60 days post inoculation and monitored for 30 days.

### Quantitation of treponemal burdens

DNAs were extracted using the DNeasy Blood and Tissue kit (Qiagen Inc., Valencia, CA), and eluted in 100-200 μl AE buffer. DNA concentrations were determined by NanoDrop^TM^ (Thermo Fisher, Pittsburgh, PA) or absorbance at 260/280 nm. qPCR of *polA* (*tp0105*) was performed as described (91).

### Molecular typing

Subtyping based on *arp* repeats and *Mse*I polymorphisms in *tprE*, *G* and *J* was performed as described (52). Strain typing was based upon sequence variability in *tp0548* described by Marra *et al*. (51) using primers listed in Table S4. Strain typing was based upon sequence variability in *tp0548* partial sequences as described.

### Nested PCR and sequencing of the β-barrel-encoding regions of *tp0558*, *tprC* (*tp0117*)*, tprD* (*tp0131*), and *bamA* (*tp0326*)

Table S3 lists unpublished primers used in this study. Nested PCR of *tp0558* was performed as described by (48). First round amplifications of the *tprC* and *tprD* β-barrel encoding regions were performed using GoTaq^®^ Flexi DNA polymerase (Promega, Madison, WI) according to manufacturer’s instructions. The resulting amplicons were gel-purified using the QIAquick Gel Extraction Kit (Qiagen) and then re-amplified using internal (2^nd^ round) primers and ExTaq (Clontech, Mountain View, CA). Nested PCR for *bamA* was carried out using GoTaq^®^ Flexi according to manufacturer’s instructions. 2^nd^ round amplicons for *tprC*, *tprD* and *bamA* were gel-purified and sequenced in both forward and reverse orientations. For patients Cali_84 and Cali_133, *bamA* 2^nd^ round amplicons also were cloned into pCR2.1-TOPO vector (Invitrogen) according to manufacturer’s instructions and ten individual clones sequenced.

### Cloning, expression, and purification of BamA L4 loops

Cloning, expression, and purification of the L4 loops of *TPA* Nichols and Mexico A was described previously (37).

### SDS-PAGE and immunoblotting

Recombinant His-tagged proteins were resolved by AnykD^TM^ Mini-Protean TGX gels (Bio-Rad) and transferred to nylon-supported nitrocellulose. Membranes were blocked and then probed overnight at 4°C with normal or immune rabbit serum at dilutions of 1:500 or with normal human or human syphilitic (Cali_84, Cali_123 and Cali_133) sera at dilutions of 1:250. Bound antibody was detected with horseradish peroxidase (HRP)-conjugated goat anti-rabbit antibody (Southern Biotech, Birmingham, AL) or HRP-conjugated goat anti-human IgG antibody (Pierce, Rockford, IL) at dilutions of 1:30,000. Immunoblots were developed using the SuperSignal West Pico chemiluminescent substrate (Thermo Fisher Scientific, Waltham, MA).

### Sequence accession numbers and phylogenetic analysis

Table S4 contains GenBank accession numbers for the *tp0558, tprC*, *tprD/D2* and *bamA* β-barrel sequences. Sequence alignments were performed using MacVector v.16.0.8 (Apex, NC). Multiple sequence alignments were performed using either MacVector or fastx_collapser from the Fastx-toolkit (version 0.0.14) (http://hannonlab.cshl.edu/fastx_toolkit/). Genes evolving under positive selection were identified with maximum likelihood method (92) implemented into PAML version 4 (93) and its user interface PAMLX (94). Site models of PAML allow the ratio of nonsynonymous/synonymous mutations (ω) to vary in each codon (site) in the gene. Branch-site models search for positive selection in lineages where different rates of ω may occur (95). Two site models and one branch-site model of PAML were used.

### Structural modelling and epitope prediction

Algorithms from the TMBpro web server (58) were used to generate the three dimensional structures for the β-barrels of TprC (Nichols) and TprD2. The structural model of BamA (Nichols) was generated as described by Luthra *et al*. (37) using the solved structure of full-length BamA (pdb ID: 3KGP) from *Neisseria gonorrhoeae* as template and the ModWeb server (https://modbase.compbio.ucsf.edu/modweb/). Discontinous epitopes were predicted using the DiscoTope 2.0 server (59), and the calculated epitopes were projected onto their respective structural models using Chimera (96). The electrostatic potential display was generated using ICM MolBrowserPro (97).

## ACKNOWLEDGEMENTS

We wish to express our gratitude to Susan Philip, San Francisco City Clinic, Population Health Division, San Francisco Department of Public Health, and Allan Pillay, Centers for Disease Control and Prevention, for providing extracted DNAs from SF patients. We are indebted to the unwavering enthusiasm and support of Nancy Saravia, CIDEIM staff members Maria Alejandra Castrillón, Juan Pablo Garcés, Luisa Rubiano and Laura Potes, The Cali Public Secretary of Health, and the health care providers from the public health hospital network who participated in patient recruitment. This work was partially supported by NIH grants R01 AI26756 (JDR), R03 TW009172 (ARC and JCS), Colciencas Grant #222956933229 (ARC), GACR grant GA17-25455S (DS), MH grant 17-31333A (DS), and by the generous support of the Department of Research, Connecticut Children’s Children Medical Center.

Portions of this work were presented at the STI & HIV World Congress, Rio de Janeiro, Brazil, 9 to 12 July 9, 2017.

The authors report no commercial or other associations that might pose a conflict of interest.

## Supplemental Material Figure Legends

**Table S1. Summary of available geographic, clinical, and demographic data for clinical samples.**

**Table S2. Distribution of clades and OMP allelic variants in *TPA* reference genomes**

**Table S3. Oligonucleotide primers used in these studies.**

**Supplemental Figure S1. Clade assignments of *TPA* clinical strains based upon *tp0548* genotypes. A.** Alignment of *tp0548* partial sequences used for genotyping. Nucleotide positions are based upon *tp0548* from SS14 (GenBank Acc. No CP004011.1). Nucleotide substitutions of G, A, T, or C are colored-coded yellow, red, green, and blue, respectively. Asterisks indicate genotypes identified in this study. Types a-i are from Marra *et al*. (1). Types k-n and p are from references (2-5). Type j belongs to *T. pallidum* subsp. *endemicum* (*TEN*) (6, 7). Type o, identified in DNA extracted from a necrophagous fly (*Stomoxys bengalensis*) trapped in northern Tanzania, aligns most closely with *TPA* Nichols (8). Type p (initially designated “o”) is based on a *TPA* isolate from Shandong, China (9). Types s, t, and x (*TPE*) were identified recently in specimens from yaws patients (10), while type u has been assigned to the *TPE* Freibourg-Blanc simian isolate (11). B. Frequency distribution and clade assignments of *tp0548* types among the 29 clinical samples in this study for which we obtained *tp0548* sequences.

**Supplemental Figure S2. Alignment of *tprC* alleles found in *TPA* reference genome sequences.** The MOSP^N^, central variable region (CVR), and MOSP^C^ β-barrel domains are shaded in blue, orange, and green, respectively. The locations of the β-barrel variable regions are indicated.

**Supplemental Figure S3. Alignment of *tprD* and *tprD2* alleles.** The MOSP^N^, central variable region (CVR), and MOSP^C^ β-barrel domains are shaded in blue, orange, and green, respectively

**Supplemental Figure S4. Alignment of *bamA* alleles based on *TPA* reference genomes.** The locations of the β-barrel variable regions are indicated.

**Supplemental Figure S6. Comparison of variable regions in β-barrel-encoding regions of *tprC* and *tprD/D2* alleles.**

